# Large-scale 3D EM connectomics dataset of mouse hippocampal area CA1

**DOI:** 10.1101/2025.04.04.647285

**Authors:** Maria Corteze, Eleonora Grasso, Mourat Vezir, Helene Schmidt

**Author notes:** Correspondence: HS.

## Abstract

The hippocampal formation is thought to be crucial for memory and learning, with subarea cornu ammonis 1 (CA1) considered to play a major role in spatial and episodic memory formation, and for evaluating the match between retrieved memories and current sensory information. While enormous progress has been made in classifying CA1 neurons based on molecular, morphological, and functional properties, and in identifying their role in behavioral tasks, a clear understanding of the underlying circuits is still missing. Here, we present the first large-scale three dimensional (3D) electron microscopy dataset of mouse CA1 at nanometer-scale resolution. The dataset is available online and can be readily used for circuit reconstructions, as demonstrated for inputs to CA1 superficial layers. Example volume segmentations show that automated reconstruction detection is feasible. Using these data, we find evidence against the long-held assumption of a homogeneous pyramidal cell population. Furthermore we find substantial possibly long-range axonal innervation of stratum-lacunosum interneurons, suggested previously to originate in L2 of MEC. These first analyses illustrate the usability of this dataset for finally clarifying the connectomic properties of mouse CA1, a key structure in mammalian brains.

## INTRODUCTION

The hippocampal formation, probably one of the most intensely studied areas in the mammalian brain, is well known for its vital role in spatial navigation and episodic memory (Scoville and Milner 1957, O’Keefe and Dostrovsky 1971, Squire 1992, Eichenbaum 2000). Subregion CA1 has been established as the key output structure of the hippocampal memory circuit. It receives processed information from CA3 and direct input from the entorhinal cortex. Functionally, area CA1 is proposed to be involved in novelty detection, amplification of the CA3 signal and input comparison (McClelland, McNaughton et al. 1995, Lisman and Otmakhova 2001, Kaifosh and Losonczy 2016, Sharif, Tayebi et al. 2021). Until recently, hippocampal computation was considered to result from random synaptic connections between a homogeneous population of pyramidal neurons (Marr 1971, Treves and Rolls 1994, Kesner and Rolls 2015), questioning how a single circuit can reliably process information under different learning conditions.

More recently it has however been recognized that CA1 pyramidal cells exhibit substantial variability, potentially enabling the formation of parallel circuit modules. Specifically, a clear morphological segregation was described along the radial axis: pyramidal neurons located closer to stratum oriens (“deep” cells) and neurons positioned closer to stratum radiatum (“superficial” cells). They were found to differ in their molecular, structural, physiological properties, afferent connections from entorhinal cortex, as well as their involvement in hippocampal oscillations (Mizuseki, Diba et al. 2011, Slomianka, Amrein et al. 2011, Lee, Marchionni et al. 2014, Stark, Roux et al. 2014, Valero, Cid et al. 2015, Danielson, Zaremba et al. 2016, Geiller, Fattahi et al. 2017, Soltesz and Losonczy 2018, Sharif, Tayebi et al. 2021, Harvey, Robinson et al. 2023). Superficial cells were found to fire more consistently during sharp wave-ripple events compared to deep cells (Stark, Roux et al. 2014). Further, a class of neurons was reported whose axon originates from a main dendrite instead of the soma (Axon-carrying-Dendrites (AcDs)). In the CA1 region, AcD cells were found to be distributed non-uniformly, preferentially clustered in the superficial layer of the stratum pyramidale compared to the deeper layers (Thome, Kelly et al. 2014). This distribution highlights the heterogeneity within the superficial and deep pyramidal cell layers of CA1 that could be accompanied by substantial local and long-distance circuit specificity.

However, a testing of such hypotheses, and the discovery of possible specificity in the connectome of this important brain area has been elusive.

Here, we report the first large-scale 3D EM dataset from mouse hippocampus CA1 that has the dimensions required for locally complete circuit analysis. In particular, it spans all hippocampal layers, extending into the neighboring dentate gyrus, has substantial width to comprise >3000 pyramidal neurons, and is of sufficient depth to contain a large fraction of the local circuitry.

With such a dataset, important questions about hippocampal circuits can be addressed. For example, the degree of neuronal variability, and circuit specificity, the existence of specific long-range inputs, and degree of synaptic specialization can be studied.

## RESULTS

We acquired a 3D-EM dataset from the hippocampal region CA1 of a P29 female mouse (Fig. 1A) using an automated tape-collecting ultramicrotome (ATUM) for ultrathin slicing at 35 nm nominal cutting thickness, followed by multibeam scanning EM image acquisition (using a 61-beam multiSEM, Zeiss at 4×4 nm^2^ in-plane voxel size and 50 ns dwell time per beam, (Hayworth, Kasthuri et al. 2006, Hayworth, Morgan et al. 2014, Eberle, Mikula et al. 2015, Sammons, Vezir et al. 2024, Sievers, Motta et al. 2024)). The total acquired image data spanned 5,086 ultrathin sections (i.e. 178 µm depth at a nominal cutting thickness of 35 nm), with an average imaged area per section of 1.6 mm^2^ (total 0.504 PB of raw data, 0.28 mm^3^ volume). From this, a first 3D EM dataset sized 1.1 × 0.92 × 0.143 mm^3^ was aligned and made available on webKnossos for online browsing, annotation and analysis (Fig. 1B, https://wklink.org/7023). This dataset spans all layers of CA1 (Fig. 1C) containing n=3,697 neurons, and extends further into the dentate gyrus. All layers of CA1 (stratum moleculare, SM; stratum lacunosum, SL; stratum radiatum, SR; stratum pyramidale, SP; and stratum oriens, SO) could be clearly delineated based on the neuropil composition at EM level (Fig. 1C).

**Figure 1.**
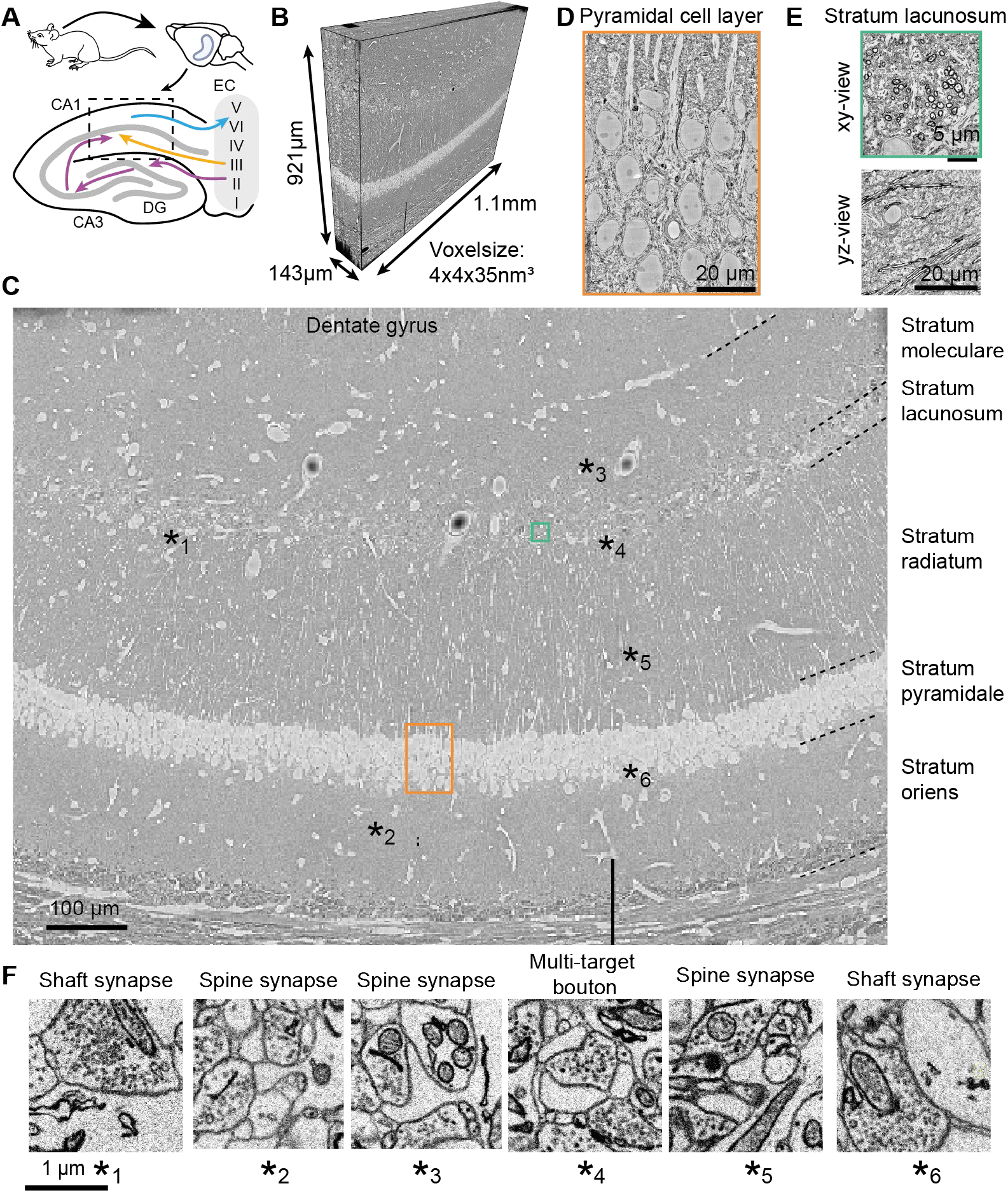
Large-scale 3D-EM dataset from mouse hippocampal region CA1. **A)** Sketch of the location of the dataset in context of the CA1-CA3-dentate gyrus (DG) anatomy of the hippocampus, and indication of so-far known long-range connections from and to the entorhinal cortex (EC). **(B)** 3D view of the outlines of the 3D-EM dataset of mouse CA1. The total imaged volume was about 1.6 mm^2^ x .178 mm, the currently aligned and browsable dataset size is 1.1 x .921 x .143 mm^3^. This 3D EM dataset is accessible on webknossos: https://wklink.org/7023. **(C)** Overview of the EM dataset in the xy plane. The detailed anatomy of CA1 is clearly discernible: the pyramidal cell (soma) layer, strata oriens, radiatum, lacunosum and separately moleculare. Note also the direct apposition of the dentate gyrus (DG). **(D)** High staining contrast and tissue integrity, also in the pyramidal cell layer, where dense somatic packing challenges fixation and staining. **(E)** xy and yz view of stratum lacunosum shows high rate of myelinated axons, and also shows high alignment quality in 3D. **(F)** Example images of synapses from various locations of the dataset, shown at high resolution. Note clarity and contrast of image data.

The tissue block had been stained using enhanced *en-bloc* staining (Hua, Laserstein et al. 2015) to provide high image contrast over the entire volume, with a special concern on the densely packed pyramidal cell layer (Fig. 1D).

Stratum lacunosum contained a high density of myelinated axons, with a clear delineation to the stratum moleculare located above (Fig. 1E). CA1 was only weakly separated from the top of the dentate gyrus (DG), raising the possibility of direct cross-innervation. Synapses onto dendritic shafts, spines, and multi-target boutons could be readily identified in the high-resolution EM data (Fig. 1F).

### Skeleton reconstructions

We next reconstructed the dendrites of neurons located in the pyramidal cell layer using skeleton reconstruction in webKnossos (https://wklink.org/7023, Fig. 2). The dataset alignment and resolution allowed the efficient reconstruction of dendritic arbors, with about 40 hours of human work load for n=20 pyramidal cells reconstructed (Fig. 2A). Deep vs. superficial pyramidal cells showed distinct morphologies and spine rates (Fig. 2B,C), suggesting high cellular variability among the pyramidal cells, also with reference to their input synapse rates. An additional rare pyramidal cell subtype, the so-called Axon-carrying-Dendrite pyramidal cell (AcD) was also found (Fig. 2D, distance of axon exit to soma, 7.1 µm).

**Figure 2.**
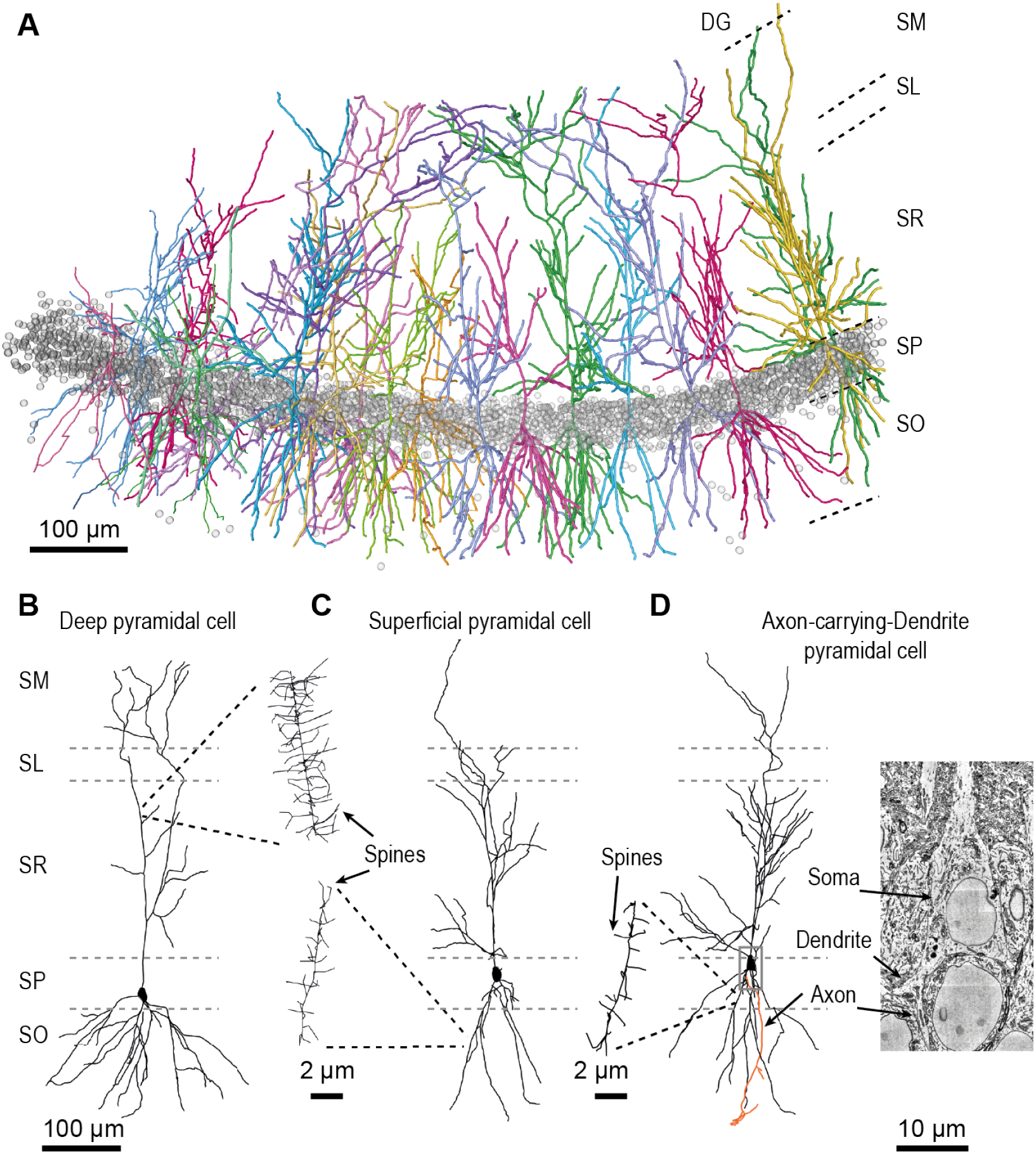
Reconstructions of CA 1 pyramidal cells. **(A)** Reconstruction of dendritic arbors of n=20 pyramidal cells using skeleton reconstruction in webKnossos (direct access: https://wklink.org/7023). Example reconstructions of deep **(B)** and superficial **(C)** pyramidal cells. Note substantial difference in dendritic morphology, and different size and rate of dendritc spines (insets). **(D)** Reconstruction of an Axon-carrying-Dendrite pyramidal cell (AcD), and EM image showing the cell body with the axon originating from the basal dendrite (right).

### Automated volume segmentation

We next investigated the usability of our large-scale dataset for dense automated volume segmentation. We used voxelytics (scalable minds, Potsdam) for image classification and segmentation on sub volumes placed in the main CA1 layers (Fig. 3). For all sub volumes, dense segmentations could be obtained that were of sufficient quality for automated neurite reconstructions (Fig. 3A-D).

**Figure 3.**
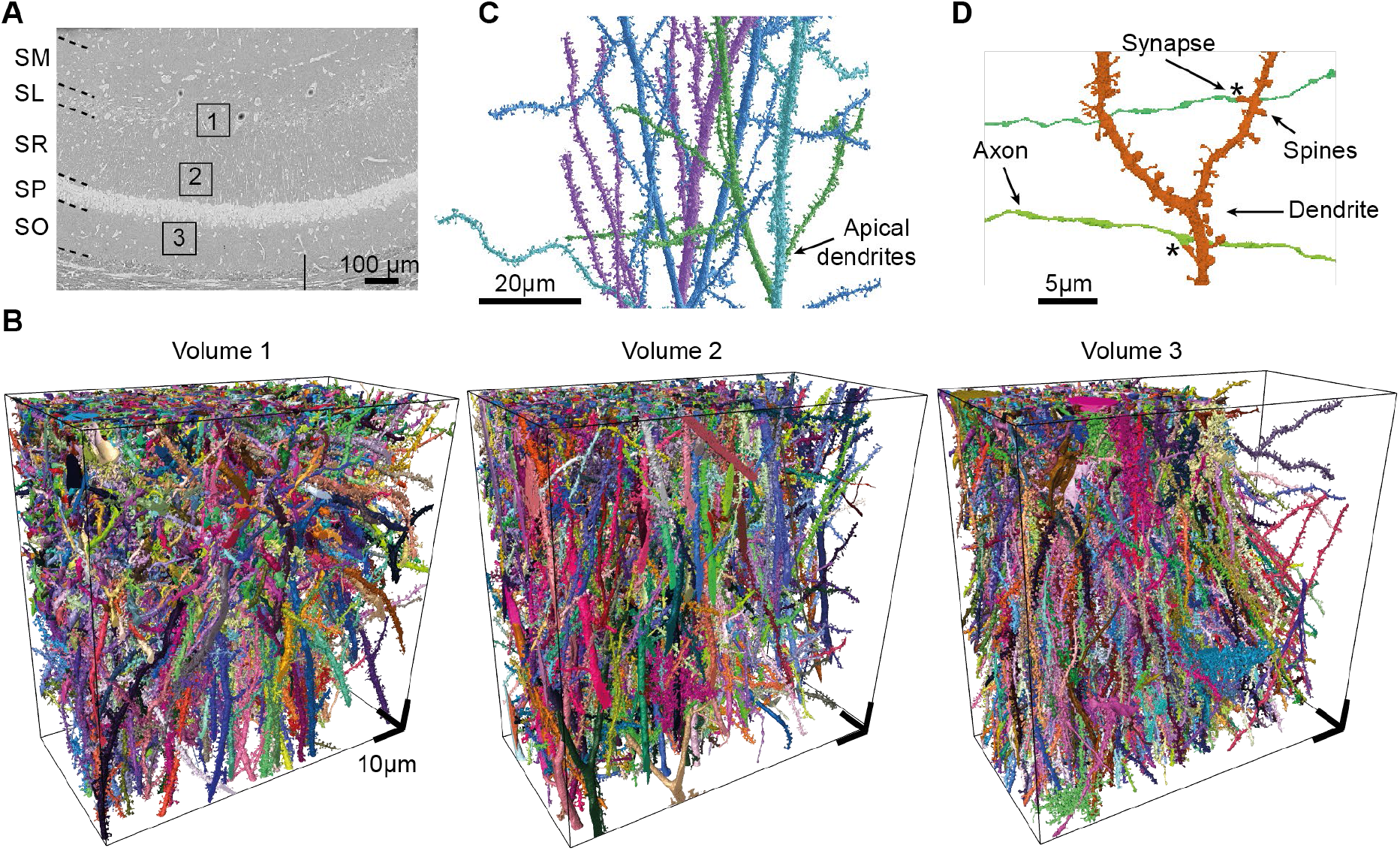
Automated volume segmentation. **(A)** Location and **(B)** examples of local (100x100x50 µm^3^) dataset volumes illustrating the quality of the automated volume segmentation of this dataset. **(C)** Example segmentation of 4 pyramidal cell apical dendrites in volume 2. **(D)** Segmentation of three axons making synapses with a dendrite of a pyramidal neuron.

### Input axons and their targets

Finally, to exemplify the usability of our dataset for analysis of target specificities of axons in superficial layers, likely long-range incoming axons, we focused on SL for finding putative long-range input axons from entorhinal cortex (Fig. 4). These input axons have previously been described to target SM and SL layers (Fig. 4A, (Steward and Scoville 1976, Suh, Rivest et al. 2011, Kitamura, Pignatelli et al. 2014). We traced myelinated axons in SL to determine whether they would branch locally and innervate targets in CA1. In fact, we found such strongly myelinated axons (Fig. 4B) that branched locally at nodes of Ranvier, making numerous output synapses. When identifying and reconstructing the synaptic targets of this axon, we found substantial innervation of smooth interneuron dendrites (Fig. 4C). The targeted interneurons were partly restricted to SL, and received as many as 29 synapses from a single axon, with clustered synaptic inputs (Fig. 4C,D). This was notable for both the substantial innervation of interneurons, the extremely high multiplicity of this innervation and the resemblance to long-range inputs from MEC layer 2 that had been reported to target interneurons in CA1 (Kitamura, Pignatelli et al. 2014). Here, we may have found the structural correlate of these substantial innervations (Fig. 4C).

**Figure 4.**
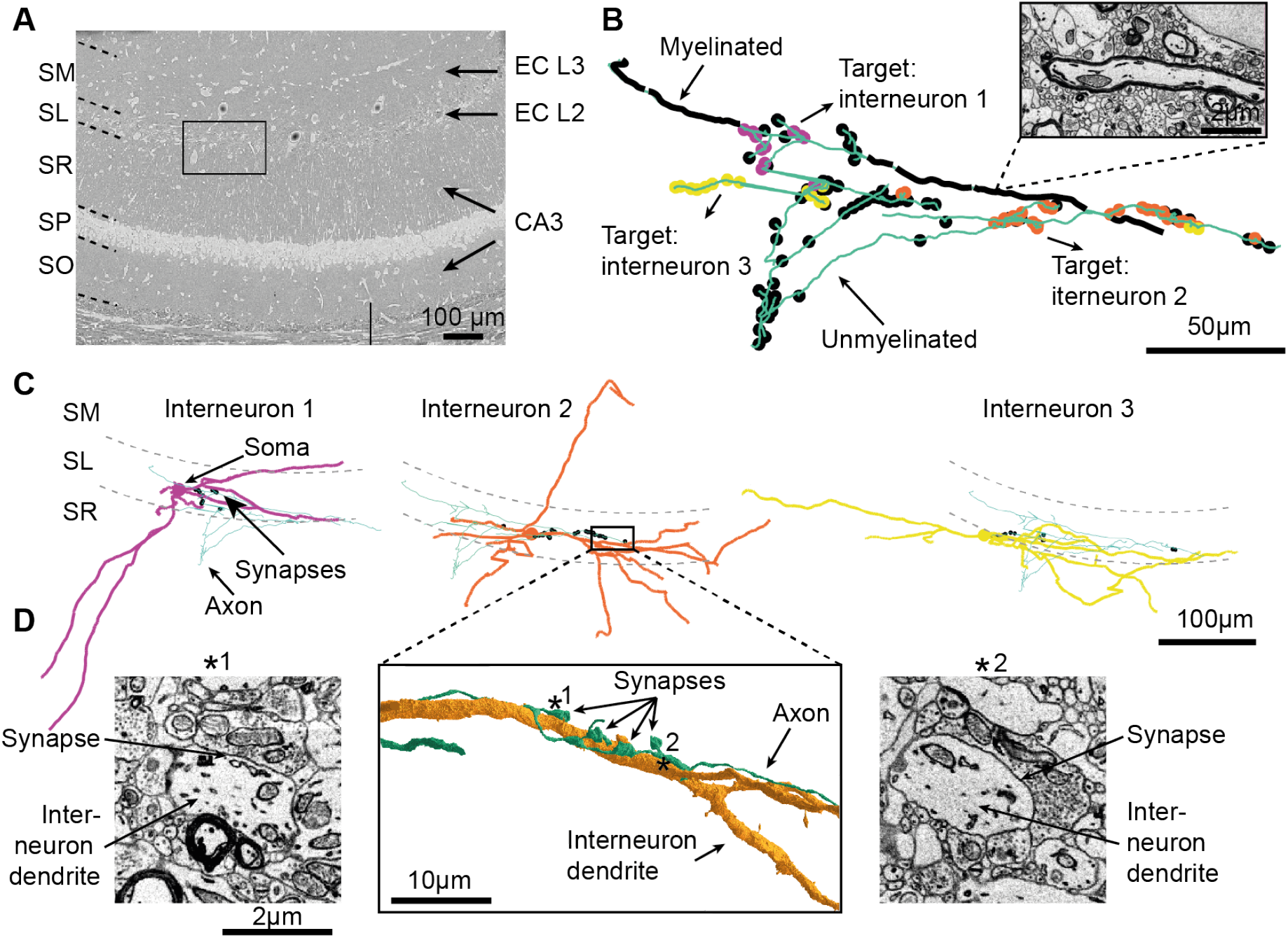
Identification of putative long-range axonal inputs to CA1 and target mapping. **(A)** Dataset overview indicating the expected long-range axonal inputs to the various layers of CA1. **(B)** Example reconstruction of a myelinated axon (green) that demyelinates and branches profusely at nodes of Ranvier. More than 29 synapses were made on interneuron dendrites. **(C)** Three example targets of the axon shown in B. The types of (LM-specific) interneurons are readily discernible. **(D)** Shows substantial synaptic clustering onto the interneuron 2 dendrite, in place to excite this interneuron from one axonal input. Location and target preference of this axon may correspond to the previously reported long-range input from MEC layer 2 (Kitamura, Pignatelli et al. 2014). Example images of the shaft synapses shown left and right.

## DISSCUSION

We obtained the first large-scale 3D EM dataset of hippocampal region CA1 in mouse, one of the most studied brain regions in mammals. Our dataset has the size, resolution, contrast, and alignment quality to readily permit cell type identification, local- and long-range circuit reconstruction, and automated volume segmentation. The data is directly openly accessible via online browsing and annotation. With this, a systematic connectomic analysis of this important brain region related to memory, association and spatial navigation can be finally conducted.

Previous EM-based synaptic analysis of hippocampal CA1 was restricted to at least one order of magnitude smaller volumes (Bartol, Bromer et al. 2015), enabling the determination of synaptic properties, but not permitting systematic circuit analysis, in particular related to pyramidal cell subtypes.

Given high image contrast and alignment quality, data analysis is immediately possible with this scale of connectomic data, as exemplified here. We expect this to be a readily useful resource for the community studying CA1 and related brain areas in mouse.

## METHODS

The experimental procedures related to tissue preparation, imaging and image registration are similar to (Sammons, Vezir et al. 2024).

### Animal experiments

All experimental procedures were performed in accordance with the law of animal experimentation issued by the German Federal Government under the supervision of local ethics committees, approved by the Regierungspräsidium Darmstadt, AZ: F126/1028, in compliance with the guidelines of the Max Planck Society.

### Tissue extraction and staining

The 3D EM dataset was acquired from a female wild-type mouse (C57BL/6J), which was born in-house and kept with littermates until weaning at P15. After weaning, the mouse was kept in a cage with their siblings (2 males and 5 females), until transcardial perfusion at P29 (RT 22 °C; relative humidity 55% (+-10%) 12-h light–dark cycle). Food and water were provided ad libitum (autoclaved water and Sniff standard mouse breeding or husbandry pellets). For transcardial perfusion, 0.1 mg/kg buprenorphine (Buprenovet, Recipharm, France) and 100 mg/kg metamizole (Metamizol WDT, WDT, Germany) were used, followed by transcardial perfusion as described in (Karimi, Odenthal et al. 2020). After perfusion, the animal’s brain was removed from the skull and kept overnight at 4°C in EM fixative made of 2.5% paraformaldehyde (Sigma), 1.25% glutaraldehyde (Serva), and 2 mM calcium chloride (Sigma) in 80 mM cacodylate buffer adjusted to pH 7.4 with an osmolarity ranging from 700 to 800 mOsmol/kg (Hua, Laserstein et al. 2015). Next, the left hemisphere was cut into 500 µm thick coronal slices using a vibratome (VT1200S Vibratome, Leica, Germany). Then, a sample spanning the whole CA1 region was extracted from the hippocampus using a 3-mm diameter biopsy punch and prepared for electron microscopy using the *en-bloc* staining method described previously (Hua, Laserstein et al. 2015, Karimi, Odenthal et al. 2020). Following dehydration, an Epon-based infiltration and embedding protocol was employed as in (Loomba, Straehle et al. 2022). In short, the sample was infiltrated sequentially with acetone and resin mixtures (ratios of 3:1, 1:1, ad 1:3) using the Epon hard mixture (5.9 g Epoxy, 2.25 g DDSA, 3.7 g NMA, 205 µl DMP; Sigma-Aldrich, USA) for 4 h, 12 h, and 4 h, respectively). The sample was then immersed in pure resin for 24 h, with replacing the resin three times (after 4 h, 12 h at 4°C, and another 4-5 h at room temperature). Finally, the sample was embedded in freshly prepared pure resin on an aluminum pin, and then the Epoxy resin was polymerized in a pre-heated oven at 60°C for 2 to 3 days (UN30pa paraffin oven, Memmert, Germany).

### Dataset acquisition

For ultrathin sectioning, the sample was trimmed into an elongated hexagonal shape (size 1.26 mm × 2.62 mm, Leica EM TRIM2, Leica Microsystems, Wetzlar, Germany). Using a custom ATUM setup, 5086 sections were collected onto plasma-treated carbon-coated Kapton tape (DuPont, coating by Fraunhofer FEP, Dresden, Germany). Cutting was performed with a diamond knife (4-mm ultra 35°, DiATOME, Nidau, Switzerland) at a nominal thickness of 35 (49 sections were acquired at 40 nm) and a cutting speed of 0.3 mm/s. After section collection, the ATUM tape was mounted onto silicon wafers (p-doped, one side polished; Science Services, Germany) with double-sided adhesive carbon tape (P77819-25, Science Services, Germany). For targeted MultiSEM imaging, overview images were obtained via light microscopy (Axio Imager.A2 Vario, Carl Zeiss Microscopy GmbH, Oberkochen). Sections were imaged using a 61-beam MultiSEM (MultiSEM, 505, Carl Zeiss Microscopy GmbH, Oberkochen) at a pixel size of 4 nm, a dwell time per pixel of 50 ns and landing energy of 1.5 kV.

### 3D image alignment

The alignment of the dataset was performed as described in (Sievers, Motta et al. 2024). First, the image transformations were computed as performed in (Scheffer, Karsh et al. 2013) and (https://github.com/billkarsh/Alignment_Projects), with custom modifications. Then, MATLAB (Mathworks, USA) based functions were used to apply the affine image transformations and convert 2D images into 3D cubes of 1,024 × 1,024 × 1,024 voxels each (https://github.com/scalableminds/webknossos-wrap). 15 sections were excluded from the alignment process due to imaging artefacts such as tissue damage. Among these, two pairs of consecutive sections were skipped, each resulting in a gap of ∼70nm. These data were then uploaded to the online annotation platform webKnossos (Boergens, Berning et al. 2017) for in-browser data visualization, neurite skeletonization, and synapse identification.

### Axon reconstructions and annotation of synapses

Axons were reconstructed by skeletonization in webKnossos (Boergens et al., 2017). Synapses were annotated as comments and visualized using custom Matlab scripts.

## Author contributions

H.S. designed research; M.C., M.V., and H.S. performed research; M.C., H.S., E.G. analyzed data; and M.C. and H.S. wrote the paper. The authors declare no competing interest.

## Acknowledgements

We thank the Connectomics Department at Max Planck Institute for Brain Research for technical support and access to high-throughput microscopy and analysis technology; in particular Smaro Soworka and Heiko Wissler for excellent technical assistance, Meike Sievers, Alessandro Motta for ATUM-mSEM and reconstruction methodology, Sahil Loomba for analysis code and Pritakshi Das for help with neurite reconstructions. Funding sources: Otto-Hahn Group funded by the Max Planck Society.

